# Seeing distinct groups where there are none: spurious patterns from between-group PCA

**DOI:** 10.1101/706101

**Authors:** Andrea Cardini, Paul O’Higgins, F. James Rohlf

**Author notes:** Corresponding Author: F. James Rohlf, e-mail address.

## Abstract

Using sampling experiments, we found that, when there are fewer groups than variables, between-groups PCA (bgPCA) may suggest surprisingly distinct differences among groups for data in which none exist. While apparently not noticed before, the reasons for this problem are easy to understand. A bgPCA captures the *g*-1 dimensions of variation among the *g* group means, but only a fraction of the ∑*n*_*i*_ − *g* dimensions of within-group variation (*n*_*i*_ are the sample sizes), when the number of variables, *p*, is greater than *g*-1. This introduces a distortion in the appearance of the bgPCA plots because the within-group variation will be underrepresented, unless the variables are sufficiently correlated so that the total variation can be accounted for with just *g*-1 dimensions. The effect is most obvious when sample sizes are small relative to the number of variables, because smaller samples spread out less, but the distortion is present even for large samples. Strong covariance among variables largely reduces the magnitude of the problem, because it effectively reduces the dimensionality of the data and thus enables a larger proportion of the within-group variation to be accounted for within the *g*-1-dimensional space of a bgPCA. The distortion will still be relevant though its strength will vary from case to case depending on the structure of the data (*p*, *g*, covariances etc.). These are important problems for a method mainly designed for the analysis of variation among groups when there are very large numbers of variables and relatively small samples. In such cases, users are likely to conclude that the groups they are comparing are much more distinct than they really are. Having many variables but just small sample sizes is a common problem in fields ranging from morphometrics (as in our examples) to molecular analyses.

## Introduction

As a general trend, modern science tends to generate a very large number of variables (*p*) from samples that can vary widely in size (*n*) and often includes few individuals relative to the number of variables. Indeed, the ‘Omics’ revolution, brought forward by the rapid advancement of informatics and molecular biology, offers some of the best examples of this trend. For instance, microarray analyses may include hundreds of genetic markers from a relatively small number of individuals (Culhane et al. 2002 is an example). However, statistically analyzing such high dimensional data with relatively small sample sizes (p/n ratios) is an important and challenging problem.

A variety of methods for dimensionality reduction are available in the statistical literature (Izenman 2008). Among these, principal component analysis (PCA) is still probably the most popular in biology. A PCA is a rigid rotation of the multidimensional space of all the variables followed by a projection of the data onto relatively few orthogonal axes that together account for as much of the overall variance as possible, though there is no reason for the axes themselves to be especially meaningful biologically. When *p* ≥ *n*, a PCA can only extract at most *n*-1 uncorrelated dimensions that, together, contain all the information about the variances and covariances of the original *p* variables (all *p* dimensions can be extracted when *p* < *n*). Often, there are dominant directions of variance so that a relatively small number of PCs may account for most of the variation. The first (higher order) PCs capture the major aspects of covariation in the sample and the later PCs the smaller ones. Bookstein (2017) first brought attention to the Marchenko-Pastur theorem that shows that large *p*/*n* ratios cause an exaggeration of the sizes of the eigenvalues for the first PCs relative to those of the last PCs, thus giving a misleading impression of the relative importance of the patterns that they seem to suggest. The initial motivation for the present paper was to investigate whether large *p*/*n* ratios might cause problems for the relatively new and increasingly popular type of PCA, between-group PCA (bgPCA). In this method a PCA is performed on the covariance matrix based on the *g* sample means (rather than on the original data matrix) followed by the projection of the original *n* samples onto these bgPC axes. Plots of these axes are then used to illustrate the distances between sample means and allow a user to judge the distinctiveness of the groups.

Phenotypic variation is complex and, although the number and choice of morphometric descriptors should be determined by the specific study hypothesis (Oxnard and O’Higgins, 2011), morphometric studies are often exploratory, tending to employ large numbers of variables, which make this discipline typically highly multivariate (Blackith and Reyment 1971). This is intrinsically true for landmark coordinate-based GM (geometric morphometrics), because each additional landmark or semilandmark adds two variables to a 2D study or three to a 3D study. While the *p*/*n* ratios are very variable (Table 1), datasets used in GM studies often have many more measurements than specimens. This is particularly common in, but not exclusive to, anthropology, the discipline in which semilandmark methods for the analysis of curves and surfaces were developed and are widely employed to study human evolution (Bookstein, 1997; Gunz and Mitteroecker, 2013; Slice, 2005). Semilandmarks are typically closely spaced sets of arbitrary points used to ‘discretize’ anatomical features, such as curves and surfaces, that are devoid of clearly corresponding landmark points; therefore, they can greatly increase the number of variables in a study. Indeed, a propensity for morphometrics to employ large numbers of variables has become especially evident in the last decade, thanks to new, cheaper and faster instruments for the acquisition and analysis of 3D images. For instance, almost 60% of about 1000 entries, retrieved at the end of 2018 in Publish or Perish (https://harzing.com/resources/publish-or-perish) using google scholar to search “geometric morphometrics AND semilandmarks”, were papers published since 2013.

**Table 1.**
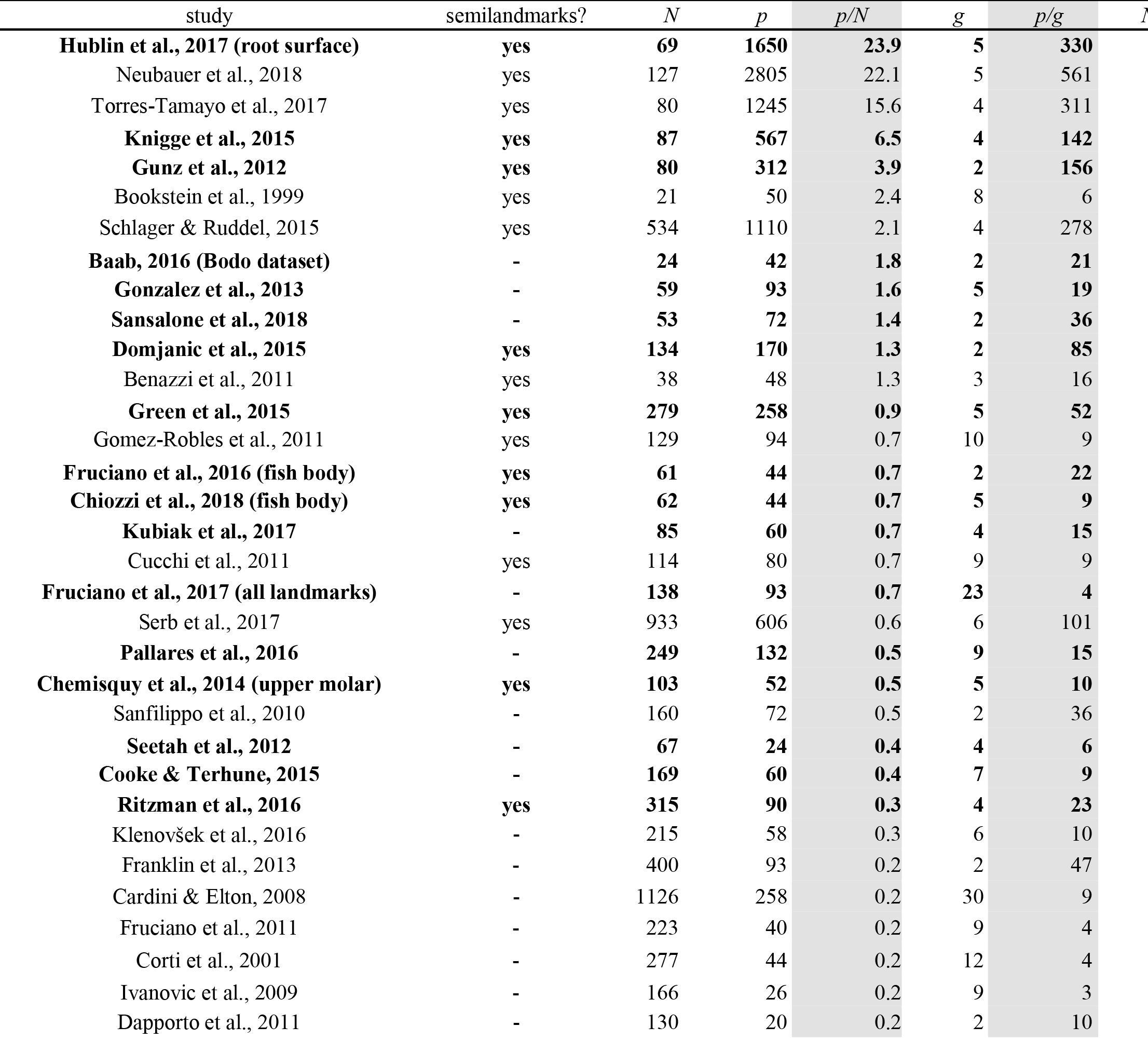
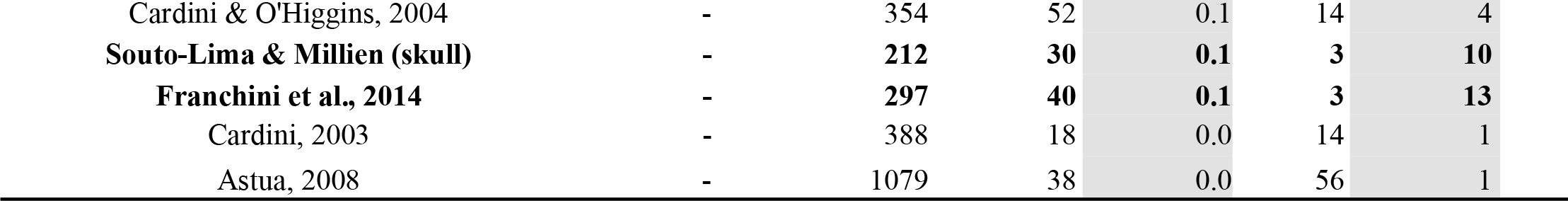
Examples of papers showing the wide range of *p/N* and *N/g* ratios used in Procrustean GM studies involving groups. The number of shape coordinates is used as a proxy for *p* (i.e., without considering the loss of dimensions in the superimposition and, if applicable, because of sliding semilandmarks or ‘symmetrization’). *N* is either the number of individuals or, if individuals were averaged in the between group analyses, the number of taxa. The average number of specimens per group (with *g* being the number of groups) is also shown. Studies using bgPCA are emphasized in bold while the columns with *pNn* and *N/g* ratios are emphasized in light grey.

| study | semilandmarks? | $N$ | $p$ | $p/N$ | $g$ | $p/g$ | $t$ |
| --- | --- | --- | --- | --- | --- | --- | --- |
| **Hublin et al., 2017 (root surface)** | **yes** | **69** | **1650** | **23.9** | **5** | **330** |  |
| Neubauer et al., 2018 | yes | 127 | 2805 | 22.1 | 5 | 561 |  |
| Torres-Tamayo et al., 2017 | yes | 80 | 1245 | 15.6 | 4 | 311 |  |
| **Knigge et al., 2015** | **yes** | **87** | **567** | **6.5** | **4** | **142** |  |
| **Gunz et al., 2012** | **yes** | **80** | **312** | **3.9** | **2** | **156** |  |
| Bookstein et al., 1999 | yes | 21 | 50 | 2.4 | 8 | 6 |  |
| Schlager & Ruedel, 2015 | yes | 534 | 1110 | 2.1 | 4 | 278 |  |
| **Baab, 2016 (Bodo dataset)** | - | **24** | **42** | **1.8** | **2** | **21** |  |
| **Gonzalez et al., 2013** | - | **59** | **93** | **1.6** | **5** | **19** |  |
| **Sansalone et al., 2018** | - | **53** | **72** | **1.4** | **2** | **36** |  |
| **Domjanic et al., 2015** | **yes** | **134** | **170** | **1.3** | **2** | **85** |  |
| Benazzi et al., 2011 | yes | 38 | 48 | 1.3 | 3 | 16 |  |
| **Green et al., 2015** | **yes** | **279** | **258** | **0.9** | **5** | **52** |  |
| Gomez-Robles et al., 2011 | yes | 129 | 94 | 0.7 | 10 | 9 |  |
| **Fruciano et al., 2016 (fish body)** | **yes** | **61** | **44** | **0.7** | **2** | **22** |  |
| **Chiozzi et al., 2018 (fish body)** | **yes** | **62** | **44** | **0.7** | **5** | **9** |  |
| **Kubiak et al., 2017** | - | **85** | **60** | **0.7** | **4** | **15** |  |
| Cucchi et al., 2011 | yes | 114 | 80 | 0.7 | 9 | 9 |  |
| **Fruciano et al., 2017 (all landmarks)** | - | **138** | **93** | **0.7** | **23** | **4** |  |
| Serb et al., 2017 | yes | 933 | 606 | 0.6 | 6 | 101 |  |
| **Pallares et al., 2016** | - | **249** | **132** | **0.5** | **9** | **15** |  |
| **Chemisquy et al., 2014 (upper molar)** | **yes** | **103** | **52** | **0.5** | **5** | **10** |  |
| Sanfilippo et al., 2010 | - | 160 | 72 | 0.5 | 2 | 36 |  |
| **Seetah et al., 2012** | - | **67** | **24** | **0.4** | **4** | **6** |  |
| **Cooke & Terhune, 2015** | - | **169** | **60** | **0.4** | **7** | **9** |  |
| **Ritzman et al., 2016** | **yes** | **315** | **90** | **0.3** | **4** | **23** |  |
| Klenovšek et al., 2016 | - | 215 | 58 | 0.3 | 6 | 10 |  |
| Franklin et al., 2013 | - | 400 | 93 | 0.2 | 2 | 47 |  |
| Cardini & Elton, 2008 | - | 1126 | 258 | 0.2 | 30 | 9 |  |
| Fruciano et al., 2011 | - | 223 | 40 | 0.2 | 9 | 4 |  |
| Corti et al., 2001 | - | 277 | 44 | 0.2 | 12 | 4 |  |
| Ivanovic et al., 2009 | - | 166 | 26 | 0.2 | 9 | 3 |  |
| Dapporto et al., 2011 | - | 130 | 20 | 0.2 | 2 | 10 |  |

|  |  |  |  |  |  |  |
| --- | --- | --- | --- | --- | --- | --- |
| Cardini & O'Higgins, 2004 | - | 354 | 52 | 0.1 | 14 | 4 |
| **Souto-Lima & Millien (skull)** | - | **212** | **30** | **0.1** | **3** | **10** |
| **Franchini et al., 2014** | - | **297** | **40** | **0.1** | **3** | **13** |
| Cardini, 2003 | - | 388 | 18 | 0.0 | 14 | 1 |
| Astua, 2008 | - | 1079 | 38 | 0.0 | 56 | 1 |

## Description of the bgPCA method

An important topic in biology is the description and interpretation of group differences in multivariate spaces Various approaches have been suggested to summarize among group variation in scatterplots (ordination methods) and to classify individuals in groups. Yet, today’s most commonly multivariate technique for separating groups is still multi-group linear discriminant analysis (DA), also known as canonical variates analysis (CVA), originally proposed by Fisher (Fisher, 1936) and Mahalanobis (Mahalanobis, 1936). However, a limit for using DA/CVA in a study is that, for statistical reliability, it requires sample sizes greatly exceeding the count of variables in the analysis (Mitteroecker and Bookstein, 2011), and indeed it is not even computationally defined if *p* > *n* − *g*. In these instances, a between-group PCA (bgPCA) has been suggested as an interesting potential alternative to explore group structure. To our knowledge, this method was originally proposed by Yendle and MacFie (1989) who called it “discriminant principal components analysis” (DPCA), though it does not involve a standardization by the within-group variation as in DA and CVA. Another early paper is Culhane et al. (2002), who applied it to the analysis of high-dimensional microarray data. While bgPCA has similarities with discriminant functions, but also, as discussed by Boulesteix (2005), has relationships to partial least-squares dimension reduction methods. Compared to DA/CVA, bgPCA is just a PCA and does not involve standardizing the variables based on the variation within groups (Seetah et al. 2012). Also, as with DA/CVA, bgPCA has been used for classification, and thus for predicting group affiliation based on bgPCs, an aim which should be achieved with a cross-validation, as exemplified by leave-one-out jack-knife used in Culhane et al. (2002) and Seetah et al. (2012). However, in contrast to a DA/CVA (Kovarovic et al. 2011; Mitteroecker and Bookstein, 2011), a bgPCA does not require *p* ≤ *n* − *g*, which is why it has been claimed that “in … between-group PCA there is NO restriction on the number of variables” (https://www.mail-archive.com/morphmet@morphometrics.org/msg05221.html).

The bgPCA procedure is used to reduce the dimensionality of multivariate data to just those dimensions necessary to account for the differences among the *g* group means. Each sample is based on *n*_*i*_ individuals for a total sample size of *n* = ∑*n*_*i*_ or *n* = *gn*_*i*_ in the case of equal sample sizes, as will be assumed here for simplicity. A bgPCA is performed by projecting the original *n*×*p* data matrix, **X**, onto the matrix, **E**, of the normalized eigenvector of the among-group SSCP matrix 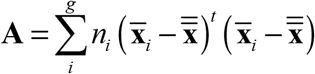, where 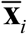 is the row vector for the mean of the *i*th group and 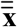 is the grand mean vector. The **A** matrix is at most of rank *g*-1 because it is a PCA of just the matrix of *g* means so only the first *g*-1 eigenvalues can be greater than zero and thus only the first *g*-1 columns of **E** need to be retained. The *n*×(*g*-1) transformed data matrix is then **X**′ = **XE**. Based on these, the transformed within-group and among-group SSCP matrices are **W**′ = **E**^*t*^**WE** and **A**′ = **E**^*t*^**AE** = Λ, the diagonal matrix of the first g-1 eigenvalues of **A** (note: the superscript “t” indicates matrix transpose; also, while the equation for **A** given above weights the mean for each group by its sample size, that may not be appropriate for many applications, see Bookstein, 2019, but it is used here for generality). Importantly, the number of bgPCs cannot be more than *g*-1. Thus, with just two groups, there are only two group means, and one needs a single dimension to represent differences between two points; thus, when *g* = 2, there is only one bgPC. If there are three groups, the differences among the three corresponding means can be fully described by a plane passing through the three mean points, and thus by just two bgPCs. With *g* > 3 the rationale is the same and the number of bgPCs is *g*-1, but the geometric representation is not as easy, because we cannot represent multivariate spaces with more than three dimensions in a single scatterplot and even a 3D scatterplot (as with *g* = 4) can be difficult to interpret (Mitteroecker et al. 2005).

## Sampling experiments

To investigate the effect of varying *p*/*n* ratios on bgPCA, sampling experiments were performed using both isotropic data (independent variables with equal means and variances called Model 1 below) and data constructed from an actual morphometric study but with no true differences among the group means (called Models 2-3 below). Fig. 1 shows the result of bgPCAs using *g* = 3 groups with the same true means (i.e., no real group differences), a constant total sample size (*n*=120), and an increasingly larger numbers of variables (*p*=12, 120 or 360). On the left (Fig. 1a) are bgPCA plots for isotropic data (Model 1, below) for *g* = 3 groups of identical size (*n*_*i*_ = *n*/3 = 40). On the right (Fig. 1b), the same *n*_*i*_, *g* and *p* are used as in Fig. 1A but based on correlated morphometric variables from real data, which have been randomly divided into three groups so that there are no real group differences. Convex hulls for each group are shown in order to identify group memberships for each sample. Rather than showing the groups superimposed as one might expect, because there are no true differences, Fig. 1 shows that bgPCA created an apparent clustering of the samples around their group means as first noticed by one of us (AC). The groups appear increasingly distinct from one another as the *p*/*n* ratio increases because larger numbers of variables are used. The effect is particularly evident for isotropic data and less pronounced but still present for correlated variables.

**Fig. 1.**
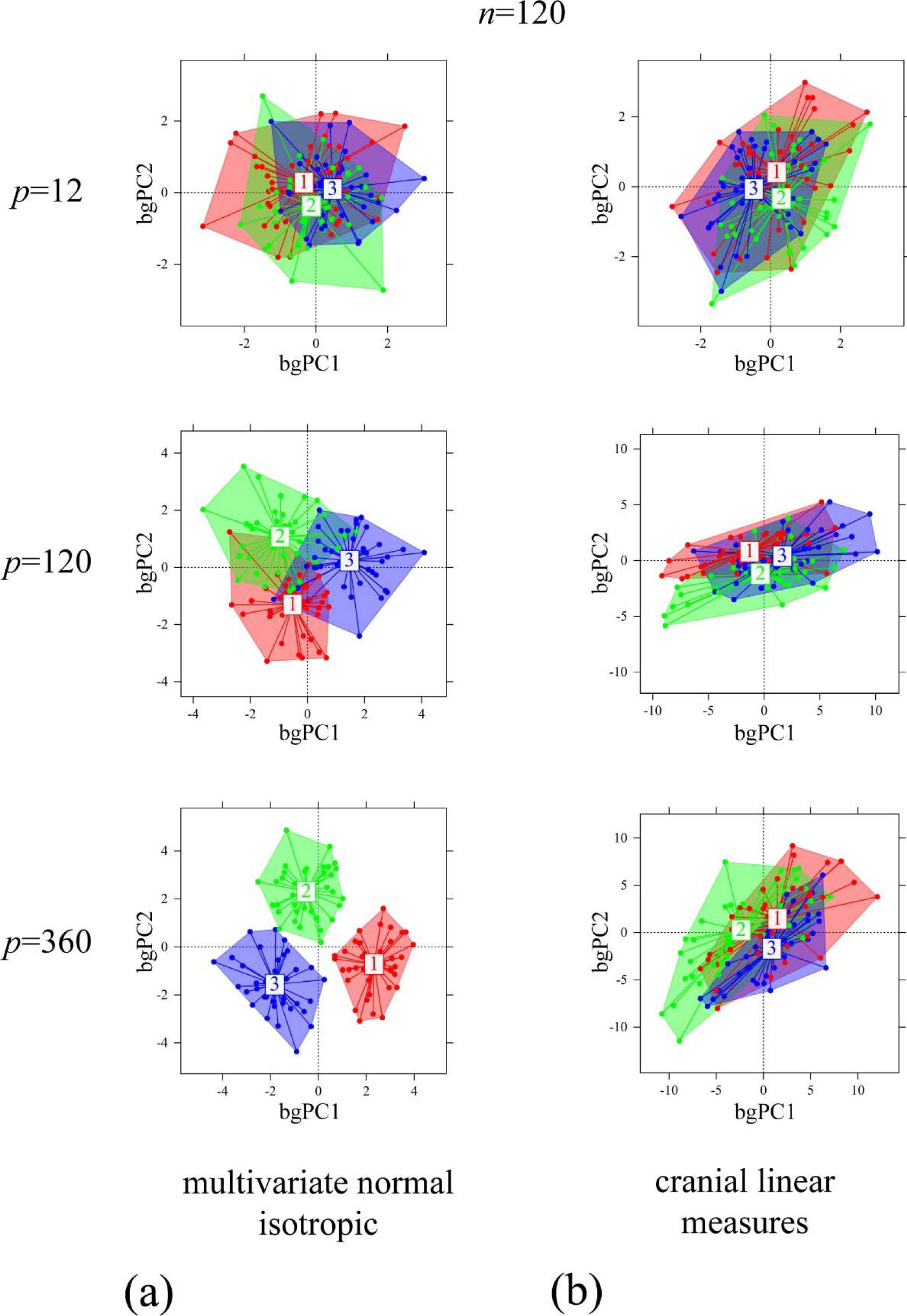
bgPCA scatterplots (computed using Morpho – Schlager, 2017 – and drawn using Adegraphics – Siberchicot et al. 2017) showing the increasing spurious separation of random groups as *p/n* increases: (a) normal multivariate isotropic (i.e., uncorrelated variables) model; (b) normal multivariate model with covarying variables (based on the covariance matrix of a set of adult male vervet cranial linear measurements).

The sampling experiments, shown in Fig. 1, were based on two different models, one (Fig. 1a) being the same as model 1 (below) and the other (Fig. 1b) being similar to models 2-3 (below). In all instances, there are no true differences among the means of the groups and the groups have the same size. Thus, in more detail, the models used in the more extensive sampling experiments described below, were:

Model 1: A purely isotropic model with p independent random normally distributed variables, each with μ = 0 and σ = 1. This model was used for Figs. 1a, 2, and 4 below.
Model 2&3: Random normally distributed variables with the same true covariance matrix as that of a real morphometric dataset, but with all means equal to zero:

Model 2: Procrustes shape coordinates from a sample of 45 adult yellow-bellied marmot (*Marmota flaviventris*) left hemimandibles. The original 2D configuration consists of 10 landmarks and 50 semilandmarks, with the semilandmarks slid in TPSRelw (Rohlf 2015) using the minimum Procrustes distance criterion. This data matrix was then used to compute the covariance matrix among the variables and its corresponding eigenvector matrix and eigenvalues. All eigenvectors that had positive eigenvalues were retained. These were then used as described below to generate random data matrices with the covariance matrix taken from the original dataset.
Model 3: Procrustes shape coordinates from a sample of 171 adult male vervet monkey skulls, which are part of a larger published dataset (Cardini and Elton 2017). There were 86 3D skull landmarks (Cardini et al. 2007; Cardini and Elton 2017). As with Model 2, as described below, these were used to generate samples of random variables with the same true covariance matrix as in the original data.

**Fig. 2.**
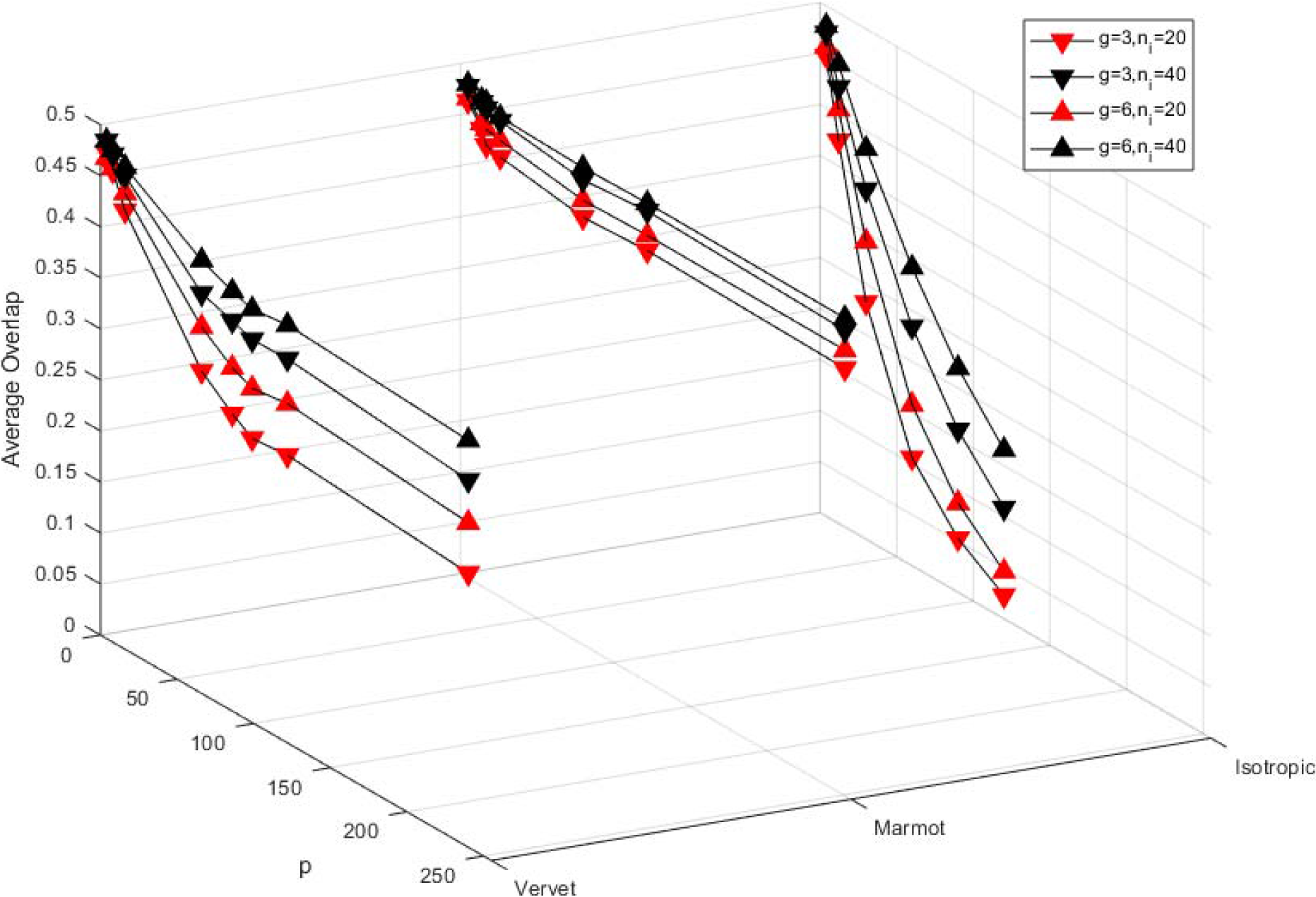
Plots of 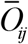 (average overlap between groups) from sampling experiments for three models: Mod1 (isotropic), Mod2 (Marmot Procrustes shape coordinates), and Mod3 (Vervet Procrustes shape coordinates), using g = 3 or 6 groups and *n*_*i*_ = 20 and 40. In all models, there is less overlap when there are fewer groups and smaller *n*_*i*_ as *p* increases.

A sample, **X**, from a population with a given true covariance matrix of ∑ was generated using the following relationship. **X** = **YEΛ**^1/2^, where **Y** is an *n*×*p* matrix of independent random normally distributed numbers with zero means and unit variances, **E** is a matrix of the *p*, *p*-dimensional normalized eigenvectors of ∑, and Λ is the *p*×*p* diagonal matrix of its eigenvalues. A difference between sampling experiments using the isotropic model (Model 1) and all others based on actual data (Models 2 and 3) is that the maximum number of eigenvectors that can be computed is limited to the number of variables in the original study because the method cannot construct more dimensions than are in the original data. For models 2 and 3, random samples of the rows (corresponding to the variables) of matrix **E** were used to generate variables. When the desired *p* was greater than the original number of variables, variables were obtained by sampling the rows of **E** with replacement.

In the sampling experiments that follow, the data were subjected to a bgPCA using code written by FJR in MATLAB and group separation was assessed by computing an index of overlap between pairs of samples. Let *O*_*ij*_ be the proportion of individuals in a group *i* that are closer to the mean of group *j*. When the dispersions in two groups *i* and *j* do not overlap, *O*_*ij*_ will be equal to 0 and will approach 0.5 for a pair of groups that overlap almost perfectly, because in that case a point is equally likely to be closest to either mean. The average, 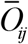 for all pairs of samples in a particular analysis is used as the measure of overlap. Initially, the amount of overlap between convex hulls was considered, but this has some unsuitable properties (such as rapid decrease in the probability of overlap as the number of dimensions increases even without the bgPCA transformation).

### What happens when *n* or *g* are changed relative to *p*?

Figure 2 summarizes the results of sampling experiments using 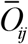 as a measure of overlap and varying *g*, *n*_*i*_, and *p*. The figure uses *n*_*i*_ rather than *n* because the total size is not relevant for the computation of average overlap as they depend on the relationships among pairs of samples and not the number of samples (and thus not on the total sample size). The sampling experiments used *g* = 3 and 6 groups, sample sizes of *n*_*i*_ = 20 and 40, and a range of values for the number of dimensions, *p*. Fig. 2 shows the expected outcome that overlap is larger when *p* is smaller, *n*_*i*_ larger, and when there are more groups. The effect of *p* is strongest for the isotropic model, but the effect is clear for all three models. The companion paper also demonstrates the effect of relaxing the assumption of equal sample sizes.

### Mathematical interpretation: why the apparent separation of groups as *p* increases?

Because the 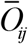 index seems difficult to work with analytically, an alternative index inspired by the partitioning of sums of squares in an anova or MANOVA was investigated for the simple null model (Model 1) used above, i.e., samples of independent normally distributed random variables from the same population. As an approximation, covariances among the variables are ignored (as they should be minimal for isotropic data) and the group differences described in terms of the traces (sums of the diagonal elements) of the usual within and among-groups sums of squares matrices, rather than the usual multivariate test statistics such as Wilks’ Lambda or Lawley-Hotelling *U* statistics, which require the computation of the matrix inversion and determinants of the sums of squares matrices.

The reader should carefully note that all expressions in Table 2 are based just on the *g*-1-dimensional space of the bgPCA transformed data. Thus, the within-group sums of squares here only refers to that part of total within group sums of squares expected in the *g*-1-dimensional subspace. This table is *not* intended for and should *never* be used for statistical testing (unlike that of a standard MANOVA, which would use the variation in the *p*-dimensional space of the original variables even if resampling procedures are used), and is specifically designed to produce an explanation for the apparent differences between groups such as shown Fig 1A.

**Table 2.**
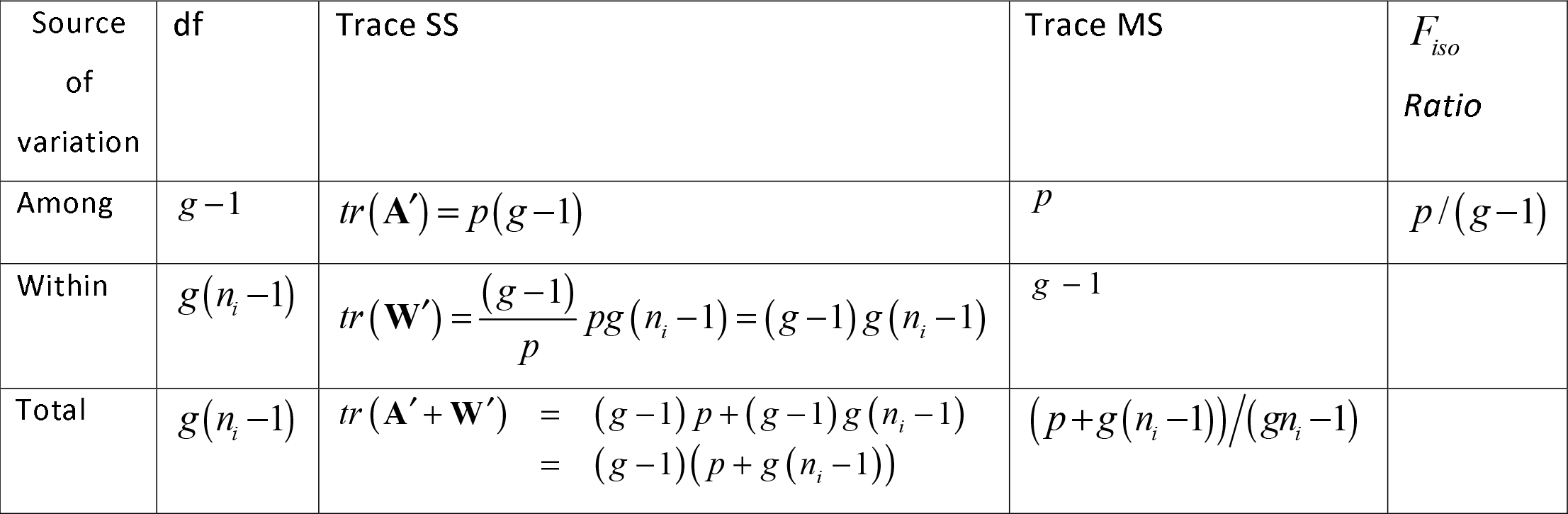
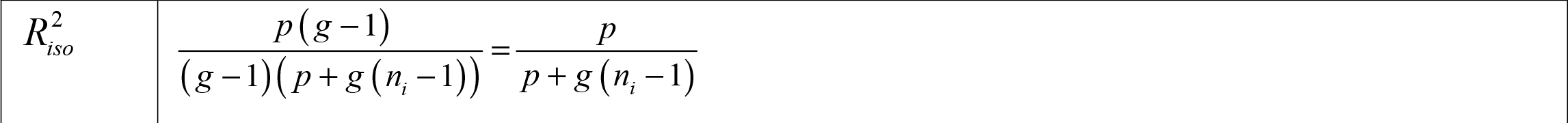
MANOVA-style table summarizing expectations after a bgPCA transformation with *g* equal-sized samples of size *n*_*i*_ all drawn from the same *p*-dimensional normally distributed population with mean **μ=0**_*p*_ (a vector of *p* zeros) and covariance matrix **∑ = I**_*p*_ (a *p*×*p* identity matrix). Because the table assumes equal-sized samples, *n*=*gn*_*i*_. The expressions for the traces of the SS matrices are given along with their MS after division by degrees of freedom. The *F*_*iso*_ ratio is also given in analogy to the usual *F* ratio and the proportion of the total variation accounted for by differences among means, 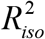, is also given. Note that these are not the usual *F* and *R*^*2*^ coefficients from an anova or a multiple regression analysis – they are expected values assuming the isotropic model, unlike a standard, MANOVA where one estimates between-group variance relative to within-group using *all* original variables, here computations are only within the *g*-1 dimensions of the bgPCA transformed data and cannot be used for statistical testing. This means that the within-group component shown in the table only refers to the residual variance left unexplained by groups in the *g*-1 dimensional bgPCA space (i.e., the within-group variation one sees in the scatterplots such as in Fig. 1).

| Source of variation | df | Trace SS | Trace MS | $F_{iso}$<br>Ratio |
| --- | --- | --- | --- | --- |
| Among | $g-1$ | $tr(\mathbf{A}') = p(g-1)$ | $p$ | $p/(g-1)$ |
| Within | $g(n_i-1)$ | $tr(\mathbf{W}') = \frac{(g-1)}{p} pg(n_i-1) = (g-1)g(n_i-1)$ | $g-1$ | |
| Total | $g(n_i-1)$ | $tr(\mathbf{A}' + \mathbf{W}') = (g-1)p + (g-1)g(n_i-1)$<br>$= (g-1)(p + g(n_i-1))$ | $(p + g(n_i-1))/(gn_i-1)$ | |

|  |  |
| --- | --- |
| $R_{iso}^2$ | $\frac{p(g-1)}{(g-1)(p+g(n_i-1))} = \frac{p}{p+g(n_i-1)}$ |

As above, let **A** represent the among-groups SSCP matrix based on all *p* variables and **E** its matrix of normalized eigenvectors. After projecting the data for all samples onto these vectors, one has a bgPCA transformed data matrix **X**′ = **XE**. At most, only the first *g*-1 columns of **E** and thus **X**′ are nonzero, so we will use only the first *g*-1 columns. Let **A**′ be the among-groups SSCP matrix based on this transformed data matrix. The sum of the eigenvalues of **A** and **A**′ are equal because all of the variation among *g* means is captured in a *g*-1-dimensional space. Similarly, one can define **W** as the within-groups SSCP matrix using the original *p* variables and **W**′ as the equivalent matrix using the projections of the data onto **E**. Note that its trace *tr*(**W**′) will, in general, be less than that of **W** because only within-group variation in the *g*-1 dimensions in which the means differ is preserved by the projection onto the *g*-1-dimensional bgPCA space. The **W** matrix has *n*−*g* degrees of freedom and thus would require min(*n* − *g*, *p*) dimensions to account for all the within-group variation.

Consider sampling experiments, such as described in the prior section for Model 1, where *n*_*i*_ specimens are in each sample (assuming equal sample size, so that *n* = *gn*_*i*_) are drawn from the same p-dimensional multivariate normal distribution, that has a mean vector **μ** = **0**_*p*_ (a vector of *p* zeroes) and a covariance matrix of ∑ = **I**_*p*_ (a *p*×*p* identity matrix). The true **W** matrix would then be (*n* − *g*) **I**_*p*_ with *tr*(**W**) ∑ *p*(*n* − *g*). The true among groups variance component matrix, ∑_*A*_, is **0**_*p*_ because there are no true differences among the population means. However, due to sampling error the expected among-groups covariance matrix is ∑+*n*_*i*_∑_*A*_. For the transformed data, the trace of the observed among-groups SSCP matrix is unchanged by the transformation because all of the variation among *g* means will be accounted for by the *g*-1 eigenvectors. However, the trace of the expected within-groups SSCP matrix will be reduced by the fraction (*g* − 1)/*p* assuming the remaining *n*−*g* dimensions of within-group variation are just a random sample of the total variation (reasonable here because, as mentioned above, there are no actual differences). These relations are conveniently summarized in the format of a MANOVA table (Table 2), but just using the trace of each matrix divided by *g*-1 as a summary of the relative amounts of within and among samples variation captured in the bgPCA space only.

Note that the *F*_*iso*_ ratio defined in Table 2 (ratio of traces of among to within group MS using only the *g*-1 bgPCs) is analogous to an *F*-ratio and is a function of just *p* and *g*. The subscript “iso” is to remind the reader that it assumes isotropic data and is not the usual *F* employed for statistical testing (that, as mentioned, should not be done using the equations of Table 2). Likewise, the “iso” in the subscript of 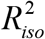 is to remind the reader that this is not the usual squared multiple correlation coefficient, because this statistic, as it is computed here using only the bgPCA variance, is only aimed at assessing the amount of group separation. Thus, a value near zero would imply that groups account for little of the total variation and values near 1 imply that most of the variation is between groups rather than within groups. Figure 3 shows plots of 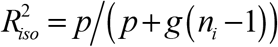 as a function of *n*_*i*_ and *p* for *g* = 3 and 6 that illustrate how 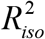 increases as a function of *p* (suggesting more distortion with more variables), but decreases as a function of *n*_*i*_ (indicating less separation of groups with larger samples). For a given *p* and *n*_*i*_, if *g* is smaller, and therefore also *n* = *gn*_*i*_ is smaller, the denominator in the 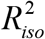 formula is reduced and 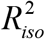 becomes larger, which is why the 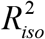 surfaces in Figure 3 are higher for *g* = 3 than for *g* = 6. This is because adding more groups increases the dimensionality of the bgPCA space and thus should account for a larger proportion of the within-group variation.

**Fig. 3.**
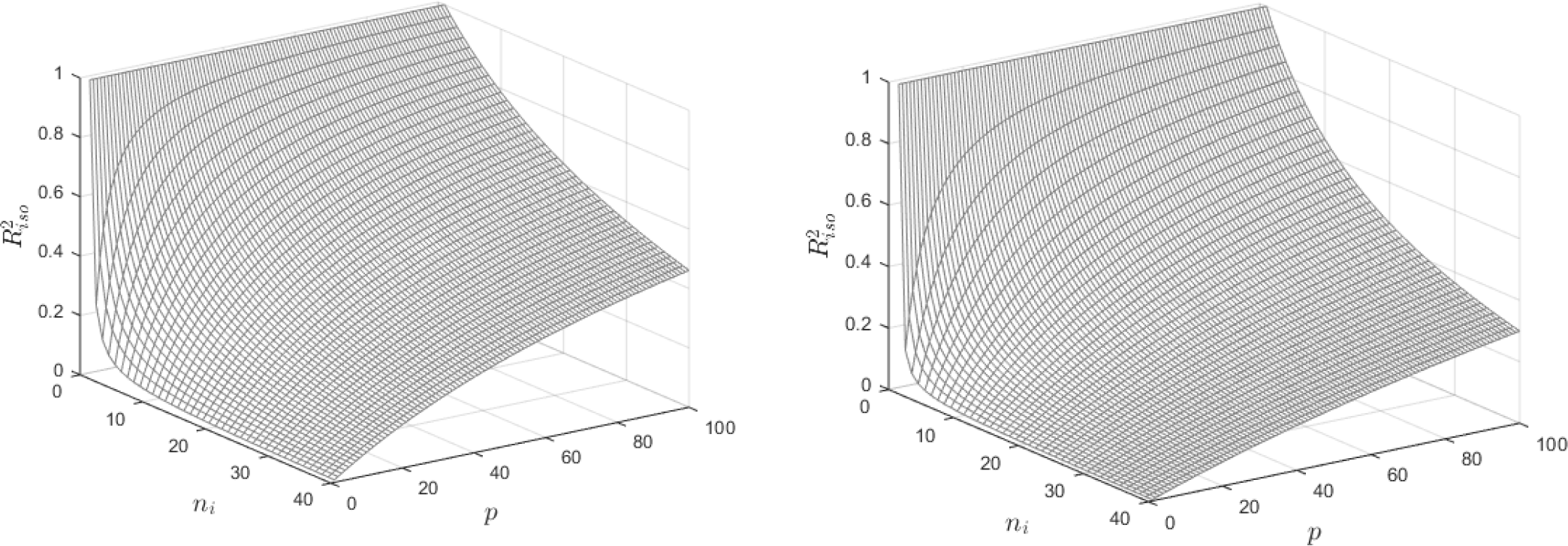
Expected relationship between 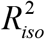 and *n*_*i*_ and *p*. A. For *g* = 3. B. For *g* = 6 groups. Note that the height of the surface is lower when larger sample sizes are larger, more groups, and fewer variables (see Table 2).

The reader should note that larger 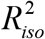 implies more separation and thus less overlap as measured by 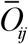. Fig. 4 shows a scatterplot 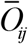 as a function of 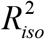 using the data from Fig. 2. The slope of the relationship differs for data from the different models. The slope is less steep for the models with correlated variables. Within each dataset the scatter corresponds to the effects of different values of *g* and *n*_*i*_. The 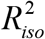 statistic is somewhat ad hoc, but Figure 4 (below) shows that it is a useful predictor of overlap for isotropic data.

**Fig. 4.**
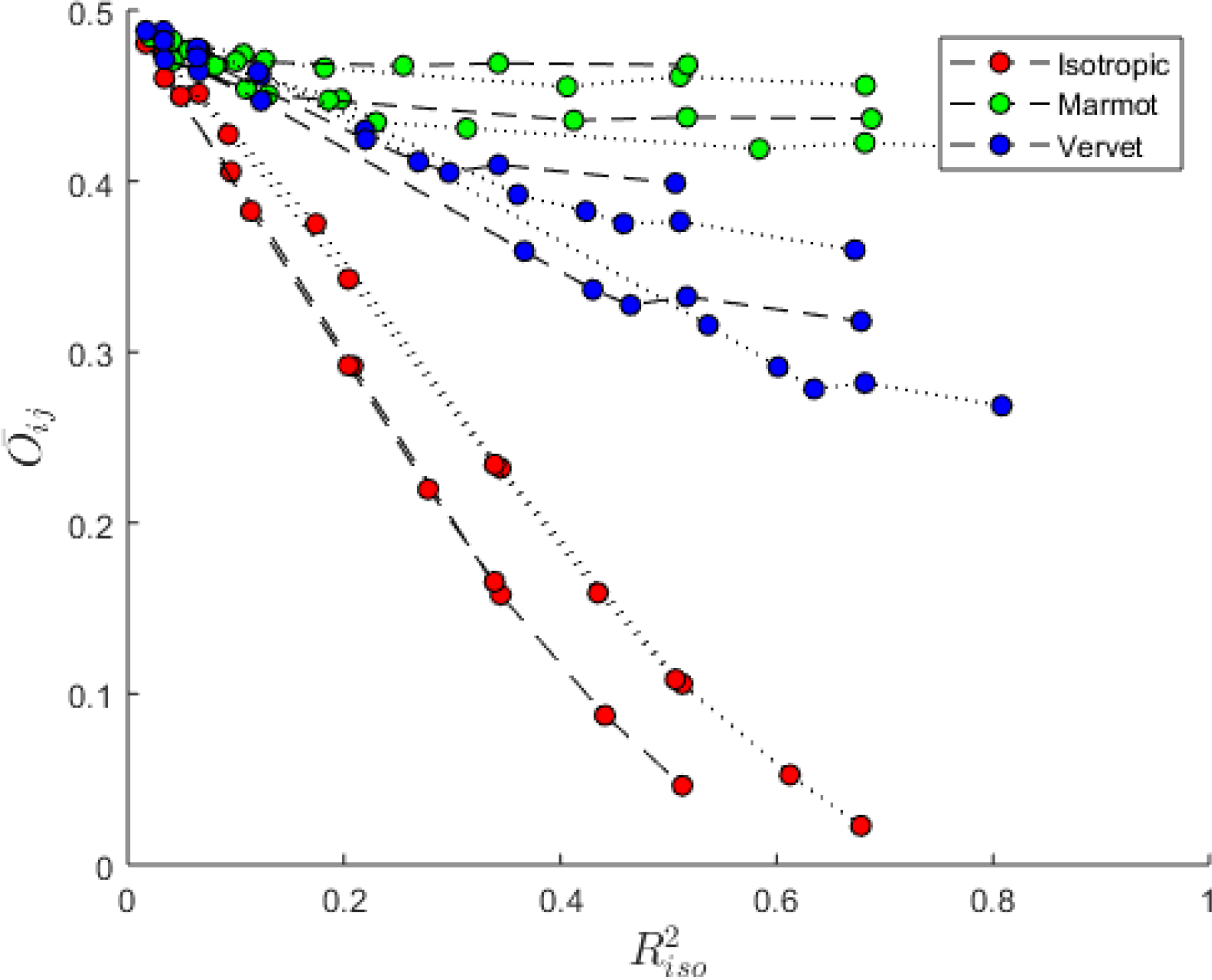
A scatterplot of 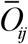 (average overlap between groups) against 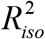 using the results of the sampling experiment shown in Fig. 2. Within each dataset it shows a tight negative relationship between 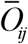 and 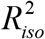 with a shallower slope for datasets that have more highly correlated variables. Dotted lines connect points for *g* = 3 groups and dashed lines for *g* = 6 groups. For isotropic data 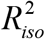 is smaller when there are more groups. Curves for different sample sizes are plotted but indistinguishable.

The expressions in Table 2 are compared in Table 3 with the results from two sampling experiments. The example in the upper half is for the case where there are fewer variables but larger sample sizes in each group. The second for the case where the number of variables is larger and sample sizes are smaller. The values are averages over 10,000 replications and show the close agreement with the expected values (given in parentheses) computed using the formulas from Table 2.

**Table 3.** Two examples of sampling experiments showing averages based on 10,000 replicates of the null model with all samples drawn from the same independent and normally distributed population with mean 0 and variance 1. Expected values based on Table 2 are given in parentheses. The upper table is an example with smaller *p* and large sample sizes. The lower table has a larger *p* and smaller sample sizes. As with Table 2, all computations are done using only using the *g*-1 = 2 dimensions from a bgPCA. Note that, unlike the formulas in Table 1, the traces are divided by *g*-1 to give an average diagonal element.

| $p = 20, g = 3, n_i = 40$ | | | | |
| --- | --- | --- | --- | --- |
| Source of variation | df | Trace SS/($g-1$) | Trace MS/($g-1$) | $F_{iso}$ Ratio |
| Among | 2 | 20.0380 (20) | 10.0190 (10) | 10.1124 (10) |
| Within | 117 | 116.9471 (117) | 0.9995 (1) |  |
| Total | 59 | 136.9851 (137) | 1.1511 (1.1513) |  |
| $R_{iso}^2 = 0.1460$ (0.1460) | | | | |

| $p = 80, g = 3, n_i = 10$ | | | | |
| --- | --- | --- | --- | --- |
| Source of variation | df | Trace SS/($g-1$) | Trace MS/($g-1$) | $F_{iso}$ Ratio |
| Among | 2 | 79.96975 (80) | 39.9849 (40) | 41.4724 (40) |
| Within | 27 | 27.0319 (27) | 1.0012 (1) |  |
| Total | 29 | 107.0016 (107) | 3.6897 (3.6897) |  |
| $R_{iso}^2 = 0.7469$ (0.7477) | | | | |

### The effect of covariation among variables

The isotropic Model 1, used in the previous section, is based on the unrealistic assumption that the variables are independent and have equal variances. Intuitively, one might expect that data with highly correlated variables might be less prone to overestimating of the degree of group separation, and indeed the sampling experiments presented in Figs. 1B, 2 and 4 do show less spurious separation for data with correlated variables (i.e., the models using vervet and marmot covariance matrices). If, as an extreme case, because of a strong correlation between variables, all of the variation in a dataset could be accounted for with just *g*-1 dimensions, then all of the within-group variation would also be captured by the *g*-1 among-groups dimensions of the bgPCA and no information would be lost. The 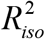 statistic described above should then be close to 0 and 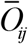 should measure the correct amount of overlap between groups, which should be close to 0.5 if there are no real groups).

In order to investigate the effect of covariation using sampling experiments, one must specify a model for the pattern and strengths of the correlations. The selection of a model can be simplified because one can rotate the data matrix to its principal axes, so that one need only consider models that differ in how the eigenvalues decrease as a function of their number. For independent variables they would decrease somewhat according to the Marchenko–Pastur formula (Bookstein 2017), but for highly correlated variables they would decrease more rapidly. A very simple model is that the logs of the eigenvalues, ln (*λ*_*i*_), decrease linearly as a function of the log of their number, that is, ln (*λ*_*i*_) = *a* − *b*ln(*i*) or as *λ*_*i*_ = *e*^−*bi*^, where *a* is a constant greater than 0 (ignored here) and *b* determines how rapidly the eigenvalues decrease. This approach also models the effect of unequal variances for the different variables. More realistic models with a factor structure could have been investigated, but this model seems sufficient to illustrate the effect of different proportions of the variance being accounted for by the first *g*-1 dimensions. Fig. 5A shows examples with *b* varied from 0 to 1. Larger values of *b* yield increasingly rapid declines of successive eigenvalues, which imply stronger correlations among variables.

**Fig. 5.**
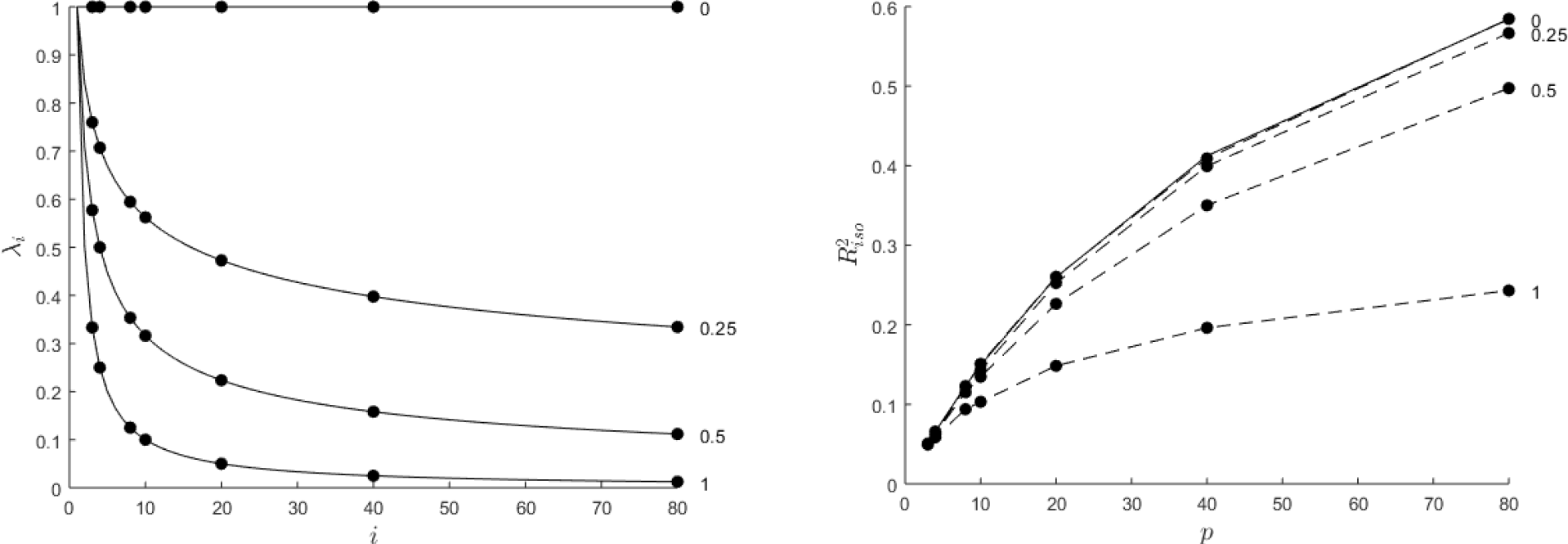
A. Plot showing the effect of varying *b* the rate of decrease of the eigenvalues (λ) for a hypothetical covariance matrix with *p* = 80 variables. The curve for *b* = 1 is similar to those usually observed in morphometric data. B. Plot showing 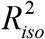 values for the results of sampling experiments for simulated data based on the models shown in Fig 5A. The slope *b* was varied from 0 to 1 to increase the level of correlation among the variables. Experiments were performed using 1000 replicates for *g* = 3 groups of size *n*_*i*_ = 20. The solid line shows the expected relationship, 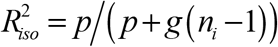, for uncorrelated data that closely matches the results from this sampling experiment. This plot shows that for the bgPCA method the proportion of the total variance accounted for by the variance among groups is expected to increase as the number of variables increases but less so as the overall level of correlation among the variables increases. For large *n*_*i*_, the slope of the curve would approach the abscissa if the correlations were such that only the first *g*-1 eigenvalues were greater than 0.

Fig. 5B shows the results of, sampling experiments with *g* = 3 groups of *n*_*i*_ = 20 observations each, with *p* ranging from 3 to 80, and each replicated 1000 times, for the b values used in Fig. 5A. The effect of increasing correlations among the variables was to reduce the size of the expected 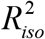 statistic implying a larger 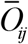 and thus less spurious clustering of points around the means. Many morphometric datasets follow patterns like that shown for *b* equal to 1 or even larger values of *b*. For instance, the curve for the marmot mandible dataset (Model 2) would be even more extreme than the curve shown for *b* = 1 The curve for the vervet data (Model 3) is less extreme. Thus, it is not surprising that Fig. 1 shows that for data with highly correlated variables there will be much less spurious group separation than that found for the isotropic model (Model 1).

## Discussion

The primary focus of the present paper is on the reasons for the apparent clustering of points around the means of arbitrary groups and predicting the magnitude of this distorted summary of group differences. In contrast, the companion paper, Bookstein (2019), examines the effect of large *p*/*n* ratios on the bgPCA method in relationship to the predictions of the Marchenko-Pastur theorem as described in Bookstein (2017), along with two other aspects of the problem: the role of variations in sample sizes of the groups, and the effect of correlations among the variables based on a variety of factor models. It also suggests ways of evaluating the impact of these effects when analyzing actual data sets. We use sampling experiments and examples from our own field, morphometrics, i.e. the quantitative study of biological forms (Blackith and Reyment 1971, Bookstein 1991). However, the issue and its implications are general and apply similarly to multivariate data used to compare groups in other fields such as genetics.

In our analyses we found that bgPCA ordinations may tend to exaggerate differences between groups relative to the amount of within-group variation. In extreme cases, with few groups, small samples and very many variables, bgPCA may consistently show perfect separation of the groups even when there are no true differences among group means. This is in part because the *g*-1 dimensions of a bgPCA capture the entire amount of variation among the *g* group means, but only a fraction of the variation within each group when *p* > *g*-1. Thus, most of the variance within groups is lost, when *p* is much larger than *g*-1. With small samples, the groups may appear quite distinct, but any apparent group differences will largely be an artefact of very large sampling error (Cardini & Elton 2007; Cardini et al. 2015). This is because any inaccuracies in group mean estimates are completely captured by the bgPCs, as if they were true differences, and used to define the *g*-1-dimensional space.

Not surprisingly, one can also see in Figure 2 that, with the same *p* and *g*, larger samples overlap more than smaller samples. Indeed, whether there are true differences or not, the range of variation within a sample is expected to increase as its sample size increases and thus there is a greater chance of overlapping.

In summary, the distortion showing a consistent spurious degree of separation between groups is not a promising property for a method that was proposed to analyze data with large numbers of variables and small samples, but the picture is complex, because the gravity of the problem, as nicely exemplified by Figure 4, varies sharply from case to case. Indeed, the severity of the distortion depends on both *g* and *n*_*i*_ relative to *p*, as well as on how strongly variables covary and whether true differences are indeed present (a case which we did not explore in our simulations). This is not unlike what Kovarovic et al. (2011) found in a study of discriminant analysis (DA). They remarked that (p. 3012): “increasing the number of predictors may increase … group separation in scatterplots of non-cross-validated DFAs, even if those predictors are random numbers which do not add any relevant information on group differences”. However, with bgPCA, this well-known problem of DA may be even more serious, because in bgPCA there is no theoretical limit to the number of variables that can be used to summarize groups and thus *p* can be much larger than *n* and *g*, as in many publications (Table 1).

Among the factors that might reduce the distortion, or even make it negligible, covariance is one of the most interesting, as it is expected in most biological datasets. The reason why covariance mitigates against the problem of bgPCA spurious group separation is that, with correlated variables, the number of independent dimensions is effectively reduced and, therefore, operationally, it is as if the *p*/*g*(*n*_*i*_ −1) ratio was smaller. The degree to which it is smaller depends on the strength of the covariances. Yet, the problem is clearly still there, as both separation and 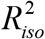 still increase with *p*. Thus, the main conclusion is the same: even with covariances, with a large *p*/*g*(*n*_*i*_ −1) ratio, not only might one see groups that appear overly separated, as in our sampling experiments, but also, if there are true groups, the differences will be inflated by a case-specific degree, which is difficult to predict a priori.

There are many reasons to expect strong covariances in studies using Procrustes-based GM. Some covariance is introduced by the fact that, for 2D data, the superimposition reduces the 2*q*-dimensional variation of the raw coordinates (with *q* being the number of landmarks) to the 2*q*−4 dimensions of shape space (Rohlf and Slice, 1990). In addition, covariation will depend on factors such as the number and distribution of the anatomical points. For example, landmarks that are very close together and closely spaced semilandmarks are expected to be highly correlated (Cardini 2018). Thus, the marmot data includes slid semilandmarks and 90% of the total variance in these data can be accounted for by just the first 10 PCs (out of the 44 possible because *n* = 45 and *p* = 120). By contrast, the vervet data requires 56 PCs (out of the 170 possible because *n* = 171 and *p* = 251) to account for the same percentage of total variance. Fig. 2 shows that the curves for the marmot data are higher (more overlap and thus less false clustering) than the curves for the vervet data (less overlap and thus stronger false separation of the groups). Note that these results do not suggest that one should purposely add highly correlated variables to reduce the distortion expected in the results of a bgPCA. Adding perfectly correlated variables to an existing dataset will not change the effective dimensionality of a dataset and thus will not alter the degree of false clustering expected in the results of a bgPCA.

On the other hand, in datasets where strong correlations among variables are expected, such as is common in GM, where additional covariance is introduced by the Procrustes superimposition itself (Rohlf and Slice, 1990) and many semilandmarks are used (because physically close semilandmarks tend to covary strongly), one might hope to circumvent some of the issues raised in this paper by reducing the number of variables used in the bgPCA. Indeed, in GM studies, it is often the case that distance matrices among specimens assessed using a few landmarks are highly correlated with those derived from the full set of landmarks plus many semilandmarks (Skinner et al, 2009; Ferretti et al. 2013; Watanabe, 2018; Galimberti et al. 2019). This can be assessed formally, for instance, through matrix correlations where testing whether full (all landmarks and semilandmarks) and reduced (a subset of the full configuration) data matrices are highly correlated. Thus, smaller ratios of *p*/*g*(*n*_*i*_ −1) can be achieved at the outset, simply by limiting the number of variables used in the study. If this is done, the resulting visualizations of shape differences among specimens will be less detailed, because fewer landmarks are used, but results of bgPCA will be less likely to be misleading.

It is important to bear in mind that scatterplots are not the only tool for assessing group differences. Results from a bgPCA should be complemented by tests of significance, as well as by cross-validated classifications of groups (e.g., Seetah et al. 2012). However, they must be performed using the full *p*-dimensional space (unlike the statistics in the ‘*ad-hoc*’ MANOVA Tables 2-3, using only bgPCs with the specific aim of assessing the magnitude of spurious group differences in the bgPCA sub-space).Wwith small samples, and/or negligible group separation *in the full data space*, group differences using all *p* variables will be non-significant, thus alerting the user that any appearance of group separation in bgPCA scatterplots should be regarded with extreme suspicion. Also, as one of the main aims in the formulation of bgPCA by Culhane et al. 2002 was classification, the results should be checked by cross-validating bgPCAs in the full data space. Finding a cross-validated accuracy only negligibly different from that expected by chance should warn users about likely distortions in the scatterplots.

In conclusion, big datasets are increasingly common, but having very many variables does not ‘counterbalance’ the effect of small *n;* it could make it worse, as shown here and in Bookstein (2019). Thus, we show that in attempting to assess group distinctiveness using bgPCA there is a potential trap, in that spurious apparent groupings may emerge in scatterplots, especially when the subspace spanned by the *g*-1 bgPCs does not adequately reflect within group variation, as is increasingly likely to happen when *p*/*n* is large and *g* is small. The appearance of spurious groups in bgPCA offers a good reminder of how a large number of descriptors might bring problems as well as benefits, with the problems sometimes potentially outweighing the benefits. Indeed, as with other methods (Hair et al. 2009; Bookstein, 2017), bgPCA provides another example of the potential perils of high dimensional data, and of the possible misuse of techniques and misinterpretation of findings, when the basic issues of sampling error and data dimensionality are not clearly borne in mind.

## Acknowledgement

We are very grateful to Jessica Grisenti, who carefully collected the marmot data for her undergraduate thesis and gave AC permission to use them. The authors appreciate the most helpful comments of Julien Claude who reviewed this paper.

## Dedications

The paper is dedicated to the memories of Nicola Saino (1961 - 2019) and Dennis Slice (1958 - 2019).

Nicola was one of the greatest Italian ethologists, Professor of Animal Behaviour at the University of Milan, and extraordinary ornithologist: AC will always remember with fondness the day Nicola introduced him, and other biology students, to the wonders of birdwatching; he will also never forget his brilliant example as a teacher and researcher; and he will greatly miss the passionate fights, with him, over methods.

Dennis Slice Professor in the Dept. of Scientific Computing, The Florida State University was an Evolutionary biologist and Ecologist who made major contributions to morphometrics through his scientific and software contributions, by maintaining and moderating the MORPHMET discussion group and by being a tireless supporter and educator of students and colleagues. For his contributions, he was awarded the Rohlf Medal for Excellence in Morphometric Methods and Applications in 2017. Not only was he a tireless advocate of his field but he was a wonderful colleague, always available and always thoughtful. We will miss him greatly as both a scientist and a colleague.

